# The effects of low-carbohydrate diets on the metabolic response to androgen-deprivation therapy in prostate cancer

**DOI:** 10.1101/2020.09.23.304360

**Authors:** Jen-Tsan Chi, Pao-Hwa Lin, Vladimir Tolstikov, Taofik Oyekunle, Gloria Cecilia Galván Alvarado, Adela Ramirez-Torres, Emily Y. Chen, Valerie Bussberg, Bo Chi, Bennett Greenwood, Rangaprasad Sarangarajan, Niven R. Narain, Michael A. Kiebish, Stephen J. Freedland

**Author notes:** Corresponding Authors: Jen-Tsan Chi, 1-919-6684759, 101 Science Drive, DUMC 3382, CIEMAS 2177A, Durham, NC 27708, Stephen J. Freedland, 1-310-423-0333 8631, W. Third St. Fourth Floor, Suite 430E, Los Angeles, CA 90048. Author contributions statement: JTC: data analysis, manuscript writing – original draft, review and editing. PHL: Conceptualization, data analysis, manuscript writing. VT, VB, NRN, BG, RS: methodology, data acquisition, statistical analysis. TO: statistical analysis. EC: methodology, project, administration, data acquisition, statistical analysis. BC: data analysis, CGA and AR: manuscript writing. MAK: Conceptualization, methodology, project administration, data acquisition, statistical analysis. SJF: Conceptualization, funding acquisitions, manuscript writing.

## Abstract

Prostate cancer (PC) is the second most lethal cancer for men and metastatic PC is treated by androgen deprivation therapy (ADT). While effective, ADT has many metabolic side effects. Previously, serum metabolome analysis showed that ADT reduced androsterone sulfate, 3-hydroxybutyric acid, acyl-carnitines while increased serum glucose. Since ADT reduced ketogenesis, we speculate that low-carbohydrate diets (LCD) may reverse many ADT-induced metabolic abnormalities in animals and humans. To test this possibility, we conducted a multi-center trial of PC patients initiating ADT randomized to no diet change (control) or LCD. We previously showed LCD led to significant weight loss, reduced fat mass, improved insulin resistance and lipid profiles. To determine whether and how LCD affects ADT-induced metabolic effects, we analyzed serum metabolites after 3-, and 6-months of ADT on LCD vs. control. We found androsterone sulfate was most consistently reduced by ADT, and was slightly further reduced by LCD. Contrastingly, LCD increased 3-hydroxybutyric acid and various acyl-carnitines, counteracting their reduction during ADT. LCD also reversed the ADT-reduced lactic acid, alanine and S-adenosyl Methionine (SAM), elevating glycolysis metabolites, amino acids and sulfur-containing metabolites. While the degree of ADT-reduced androsterone was strongly correlated with glucose and indole-3-carboxaldehyde, LCD disrupted such correlation. However, many LCD-induced changes were seen at 3-but not 6-month, suggesting metabolic adaption. Together, LCD significantly reversed many ADT-induced metabolic changes while slightly enhancing androgen reduction. Future research is needed to confirm these findings and determine whether LCD can mitigate ADT-linked comorbidities and possibly delaying disease progression by further lowering androgens.

**Statement of translational relevance:** Prostate cancer (PC) is the most common non-skin cancer and second leading cause of cancer-related death in men. While androgen deprivation therapy (ADT) is the main treatment for metastatic PC, it has many metabolic side effects. Previous serum metabolome analysis of PC patients receiving ADT identified reduced ketogenesis. Therefore, low-carbohydrate diets (LCD), ketogenic in nature, may reverse many ADT-induced metabolic abnormalities. We conducted a 6-month multi-center trial of no diet change (control) vs. LCD in men initiating ADT. We found that LCD reversed many ADT-induced metabolic abnormalities while slightly further reducing androgen levels. Also, LCD disrupted the diabetogenic effects of ADT, but some effects seen at 3-month were lost at 6-month, suggesting metabolic adaptation to LCD. These data suggest the potential metabolic benefits of LCD with potential to enhance ADT efficacy. Larger studies testing whether LCDs mitigate metabolic side effects and slow disease progression are warranted given acceptable safety profiles, metabolic benefits, and possibly lowered androgens.

## Introduction

Prostate cancer (PC) is the most common non-skin cancer and the second leading cause of cancer mortality for men in the US (1). In 2020, 191,930 new cases of PC and 33,330 deaths from PC were estimated (2). For men with non-metastatic disease who require treatment, local therapy is not always curative. While systemic therapies, such as chemotherapy and novel hormonal agents, that improve overall survival in late-stage PC (3,4), these therapies can have significant side effects including elevated risk for cardiovascular diseases (5). Moreover, the backbone of systemic treatment for PC remains androgen deprivation therapy (ADT) by decreasing male hormone levels to reduce tumor growth.

In a typical Western diet, usually 40-60% kcals come from carbohydrates, whereas low-carbohydrate diets (LCDs) usually include <20% carbohydrate kcals (6). LCDs and ketogenic diets (a very low carbohydrate diet) improve diabetes and aid with weight loss (7-9). Since the common side effects of ADT include weight gain, body fat gain, increased adiposity and insulin resistance, LCDs may benefit PC patients by minimizing the side effects during this therapy. Thus, the main objective of the CAPS1 (Carbohydrate and Prostate Study 1) study was to compare the impact of a 6-month LCD intervention vs. a control arm on the insulin resistance in PC patients initiating ADT. Results from the comparison between two arms (10) showed that at 3 months of intervention, LCD group significantly reduced weight, improved insulin resistance, hemoglobin A1c, high-density lipoprotein (HDL), and triglyceride levels. However, at 6 months, only weight loss and HDL remained significantly different between arms. We also observed other markers that reversed toward baseline at 6-month, though these were not significant in our small sample (n=29), and as such, these potential benefits of LCD deserve further examination.

Metabolomic analysis allow an integrated read-out of all the upstream biochemical pathways associated with various oncogenic genetic drivers, therapeutic intervention and lifestyle choices (11,12). Thus, metabolomic results can provide insights into various factors during tumor progression, invasion and metastasis (13). For example, the metabolomic analysis of prostate tumors identified the relevant role sarcosine-related pathways in the PC oncogenesis (14,15). Other analyses aimed to identify therapeutic targets to treat PC (16-18) or elucidate the metabolic response to the PC treatments. While ADT is the standard treatment for metastatic PC with significant metabolic side effects, however, to date, only three studies have analyzed the metabolome on serum from PC patients receiving ADT (19-21). Briefly, Saylor et al. reported 56 significantly altered metabolites after receiving 3 months of ADT, including androgen steroids, bile acid metabolites and lipid β-and Ω-oxidation (20). The second study identified seven ADT-associated metabolites, which reverted to control levels after ADT was stopped (19). Recently, we reported the serum metabolomic and lipidomic profiling of PC patients in the control arm of the CAPS1 trial after 3 and 6 months of ADT treatment (21). As expected, ADT lowered androsterone sulfate and the degree of reduction was significantly correlated with increased serum glucose, supporting a diabetogenic role of ADT. ADT also reduced 3-hydroxybutyric acid, many acyl-carnitines and indole-3-carboxaldehyde, a tryptophan-derived metabolite of the gut microbiome that serves as an agonist for the aryl hydrocarbon receptor to regulate mucosal immunity (22). Given the metabolic benefits observed in patients in the LCD arm of the CAPS1 study (10), the main purpose of this secondary analysis is to evaluate the metabolic impact of the LCD intervention in these PC patients during ADT using serum metabolomic technology. In addition, we will evaluate how LCD affects the efficacy of ADT and potential therapeutic benefits.

## Participants and Methods

### Study Design

As previously described (10) (21), IRB approval was obtained at Duke, Durham Veterans Affairs Medical Center [VAMC], and Greater Los Angeles VAMC. PC men initiating ADT were approached and if they agreed to participate, they signed a written consent that allowed future analyses. Clinical data and fasting serum samples were collected at baseline prior to ADT (BL) and three (M3) and six months (M6) post-randomization and post initiation of ADT. Key eligibility included men about to begin hormonal therapy (LHRH agonist, LHRH-antagonist, or orchiectomy) for PC with an anticipated duration of ≥6 months, BMI ≥24 kg/m^2^, and phone access to speak with the dietitian. Key exclusion criteria included medication-controlled diabetes, taking any medications that may interfere with insulin metabolism, already consuming an LCD, being vegetarian/vegan, or baseline screening HbA1c >7%. A total of 42 eligible participants enrolled and were randomized immediately after the baseline assessment. A total of 40 participants completed the baseline visit (N=19 LCD, N=21 control). Participants in the control arm received standard care and were asked to make no changes in diet or exercise pattern. Those in the LCD arm received individual coaching from the study dietitian to reduce carbohydrate intake to ≤20 gram/day and to walk for ≥30 minutes/day for 5 days/week. The study dietitian talked to the participants weekly for 3 months then bi-weekly for another 3 months. In this manuscript, we report the results of how LCD affected serum metabolomics during ADT.

### Data Collection and analysis

At each visit, weight (without shoes and in light clothing) and height were measured, and fasting blood was collected for blood chemistry. Fasting blood was analyzed for routine blood chemistry, including PSA, insulin, glucose, lipids, and high sensitivity C-reactive protein (hsCRP). PSA, glucose and lipids were measured by commercial laboratories (LabCorp for Duke and Durham VAMC sites and Greater LA VAMC clinical lab for the LA site). Insulin was measured in an electro-chemiluminescent immunoassay using an SI-2400 imager and assay kits from Meso Scale Discovery (Rockville, MD) by Duke Immunoassay Laboratory. Insulin resistance, as estimated by the homeostatic model assessment (HOMA), was calculated using the approximation (glucose*insulin)/22.5.

Fasting blood samples collected at BL, M3, and M6 were used for metabolomics analysis utilizing GC/MS-TOF (Gas chromatography–Mass Spectrometry Time of Flight analyzer), QqQ LC (Liquid Chromatography)-HILIC (Hydrophilic interaction chromatography)-MS/MS, and TripleTOF LC-RP-MS as described previously (21,23).

### Statistical Analysis

Significant changes were examined by ANOVA and shown in heatmaps and box plots. Impacted metabolic pathways were selected and mapped by using MetaboAnalyst 4.0 (https://www.metaboanalyst.ca/) (24). Pearson correlation was conducted to examine associations between clinical variables and metabolites. Volcano plots were used to visually examine the changes in metabolites at M3 and M6 from baseline. Zero transformation was performed using the attached Python scripts (supplemental file 1) modified from previous studies (25,26) to derive the ADT-induced changes of all metabolites at M3 and M6 from corresponding BL in the control and LCD arms.

## Results

### Serum metabolites and metabolic pathways altered after 3 and 6 months of ADT in the LCD arm

We measured and identified metabolites that were altered in the LCD arm between BL (immediately before ADT) and after 3 (M3) or 6 months (M6) of ADT. We identified 49 metabolites at M3 showing at least a 1.2-fold change and t-tests p<0.1 relative to BL. The top metabolites at M3, as shown in volcano plots (Fig 1A, Supplemental Table 1), included an increase in 3-hydroxybutyric acid, 3-hydroxy-3-methylglutaryl-carnitine and hydroxy-butyryl-carnitine. Also, there was a reduction in the androsterone sulfate, and glycolysis-related metabolites, including sucrose, lactic acids and fructose-6-phosphate. As several selected metabolites belong to similar metabolic pathways, we employed <u>M</u>etabolite <u>S</u>et <u>E</u>nrichment <u>A</u>nalysis (MSEA) (27) to evaluate the potential enrichment of functionally related metabolites covering various metabolic pathways. The top MSEA-identified altered pathways at M3 included alanine metabolism, citric acid cycle, estrone metabolism, carnitine synthesis, gluconeogenesis, androgen/estrogen metabolism and glycolysis (Fig 1B).

**Figure 1:**
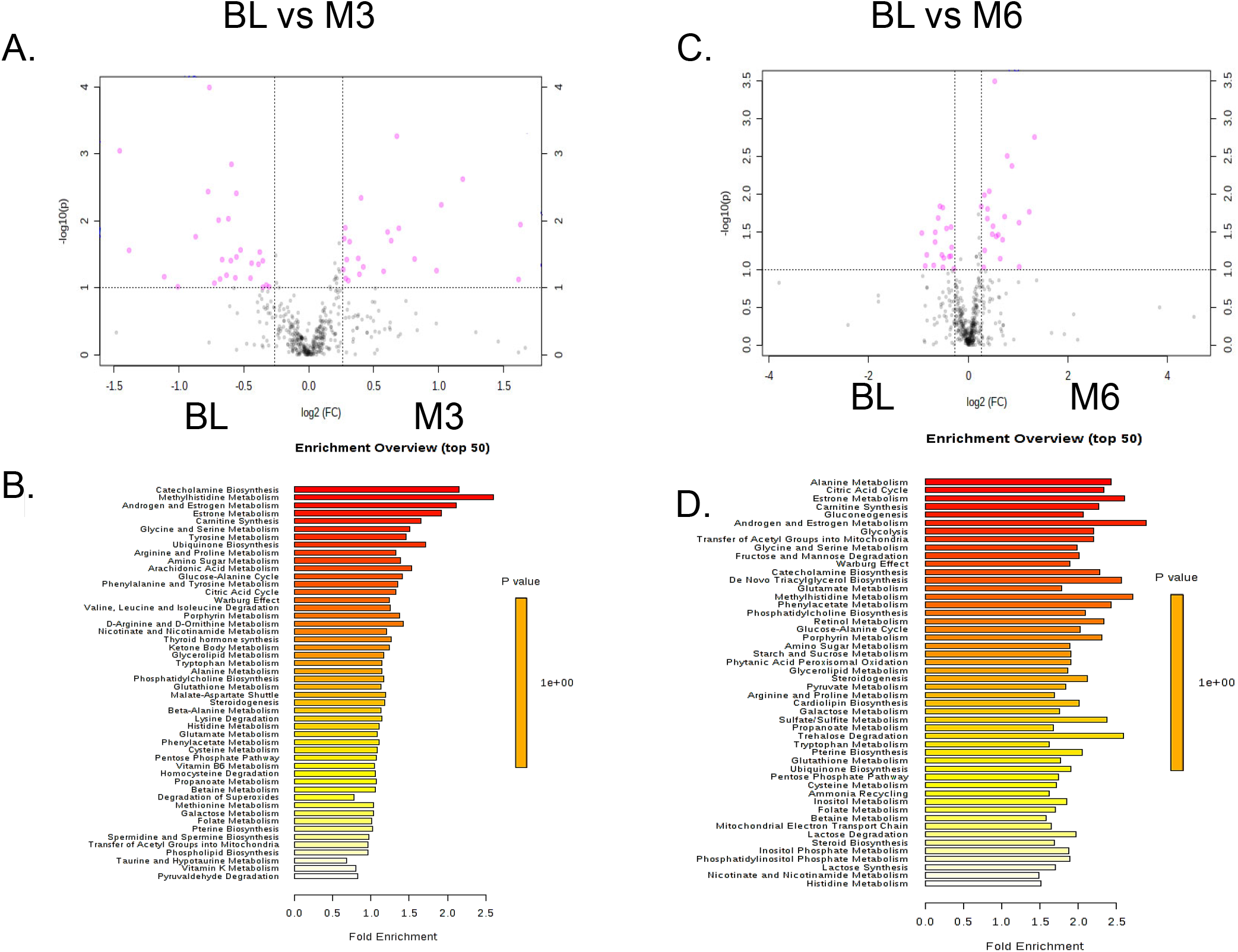
Top serum metabolites and metabolic pathways altered at 3- and 6-month after initiating ADT treatments in participants receiving LCD. (A) Metabolites were selected by Volcano plot analysis of pairwise comparison between the BL vs. M3 (A) and BL vs. M6 (C) ADT. (B, D) Top enriched metabolic pathways affected by ADT after M3 (B) and M6 (D) of LCD.

Using similar approaches, 39 metabolites were identified between M6 vs. BL as having at least 1.2-fold change and t-tests p<0.1 and shown in volcano plots (Fig 1C, Supplemental Table 2). At M6, the top changes included the reduction in androsterone sulfate, proline, sucrose and fructose-6-phosphate. Several carnitines were also found to be reduced, including dithio-acetyl-carnitine, malonyl-carnitine, and heptanoyl-carnitine. The top altered pathways at M6 identified by MSEA included catecholamine biosynthesis, methyl histidine metabolism, androgen/estrogen metabolism, estrone metabolism and carnitine metabolism (Fig 1D).

### Overview of the ADT-induced metabolomic changes in the control and LCD arms

To further understand the changes of the metabolites in response to ADT between the control and LCD arms (Fig 2A), we performed zero transformation (detailed in methods modified from (26,28,29)) to calculate the changes of all metabolites at M3 and M6 when normalized against the same metabolites at baseline (BL) in the same participant. These changes were then arranged by hierarchical clustering (Fig 2A). While there were marked individual variations in the metabolic changes among different participants, some consistent patterns emerged. First, there was a cluster of metabolites consistently reduced by ADT in most participants in both arms (Fig 2A, green bar expanded into Fig 2B). This cluster contained androsterone sulfate and other androgens, including DHEA sulfate and pregnenolone sulfate (Fig 2B). The reduction of these androgen-related metabolites is the expected metabolic effects of ADT, confirming its therapeutic effects, compliance and our analytic approaches. Interestingly, while all these androgens were reduced by ADT, androsterone sulfate was the most consistently reduced metabolite and the only androgen selected by the univariate statistics (Fig 1A, C). In addition, we noted a cluster of metabolites which were repressed in the control arm but elevated in the LCD arm (Fig 2A, red bar expanded into Fig 2C). These metabolites, affected by ADT in opposite direction in the control vs. LCD arm, included 3-hydroxyburtyric acid, hydroxybutyryl-carnitine, 2-hydroxyburtyric acid and many acyl-carnitines previously noted to be repressed by ADT in the control arm (21). These results indicate that LCD reversed the ADT-reduced ketones and acyl-carnitines levels (Fig 2C) and linking these two classes of metabolites found to be reduced by ADT(21). Therefore, some of the ADT-induced changes seen in the control persisted in the LCD arm (such as reduced androgens) while others (such as reduced ketones and acyl-carnitines) were mitigated/reversed by the LCD.

**Figure 2:**
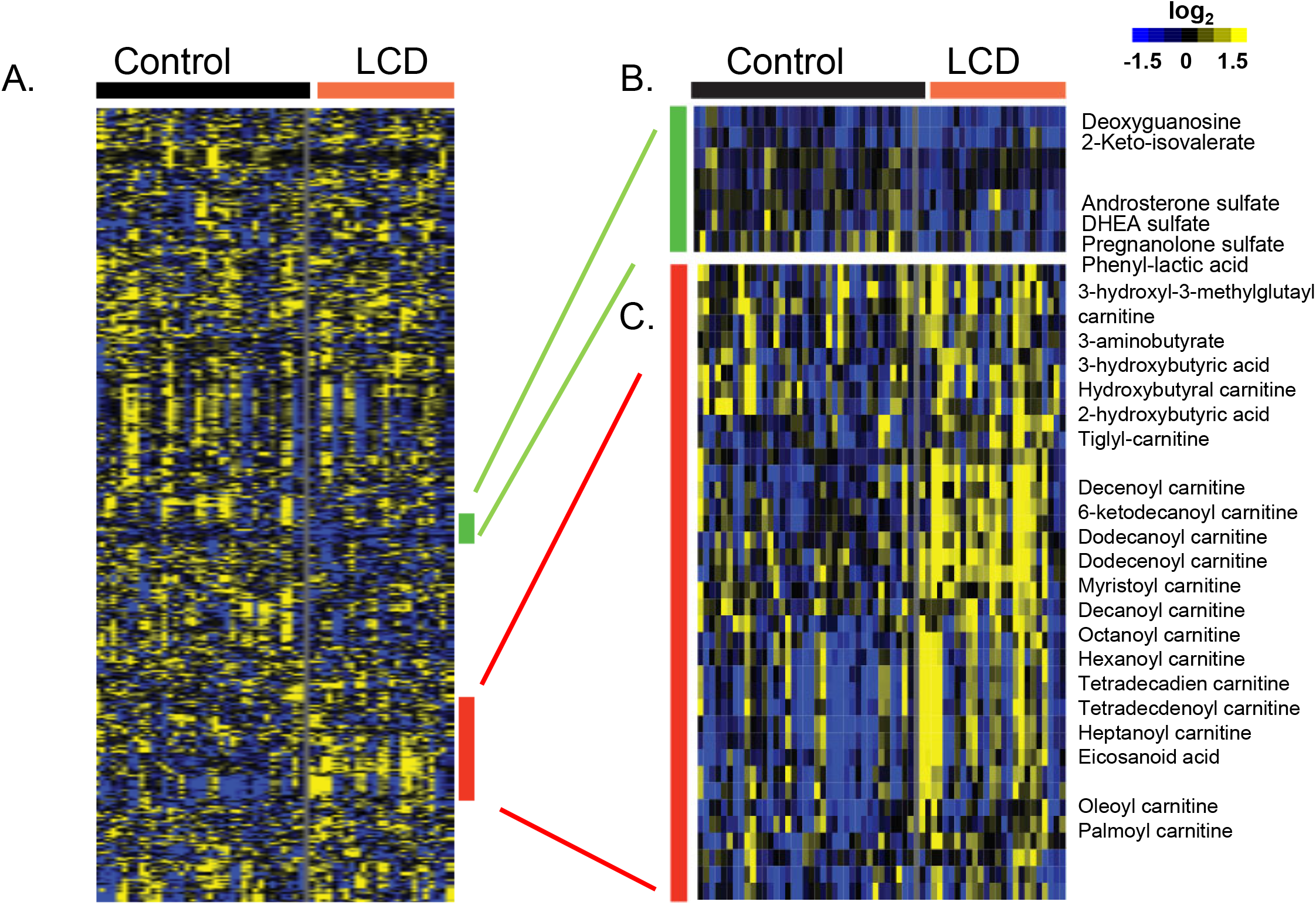
Heatmap of the ADT-induced changes in all metabolites in control and LCD arms. (A) The changes of all metabolites from BL in response to ADT were derived by zero-transformation and arranged by hierarchical clustering. (B) Yellow indicates an increase, blue indicates a reduction and black indicates no change. (B) Clusters of metabolites that were repressed (C) from BL in all samples were expanded with the names of the metabolites shown. (C) Clusters of metabolites that were repressed in control (D) samples but elevated in LCD were expanded and with the names of the metabolites shown.

### ADT-induced metabolic changes in both control and LCD arms

To identify metabolites whose changes were statistically significant between treatment groups in both arms, we applied ANOVA - Simultaneous Component Analysis (ASCA module, MetaboAnlyst 4.0) to uncover the major patterns associated with the time points and the treatments. We found that androsterone sulfate was reduced by ADT in both control and LCD arms (Fig 3A). This suggests that LCD did not affect the ADT-reduced androsterone sulfate. Instead, the degree of androsterone sulfate reduction (Fig 3B), despite not being statistically significant (M3: p=0.07, M6: p=0.08), was greater in the LCD arm than the control arm. In the heatmap (Fig 3C) showing the ADT-induced metabolic changes from BL, we noted that the reduced androsterone sulfate was co-clustered with similar reduction of other androgens, including DHEA sulfate and pregnenolone sulfate, as well as phenyl-lactic acid and phenyl-pyruvate. The ADT-reduced DHEA and pregnenolone sulfate also trended lower in the LCD arm, without reaching statistical significance. Overall, this finding suggests the potential of LCD to slightly enhance the ADT-induced reduction in androgens.

**Figure 3:**
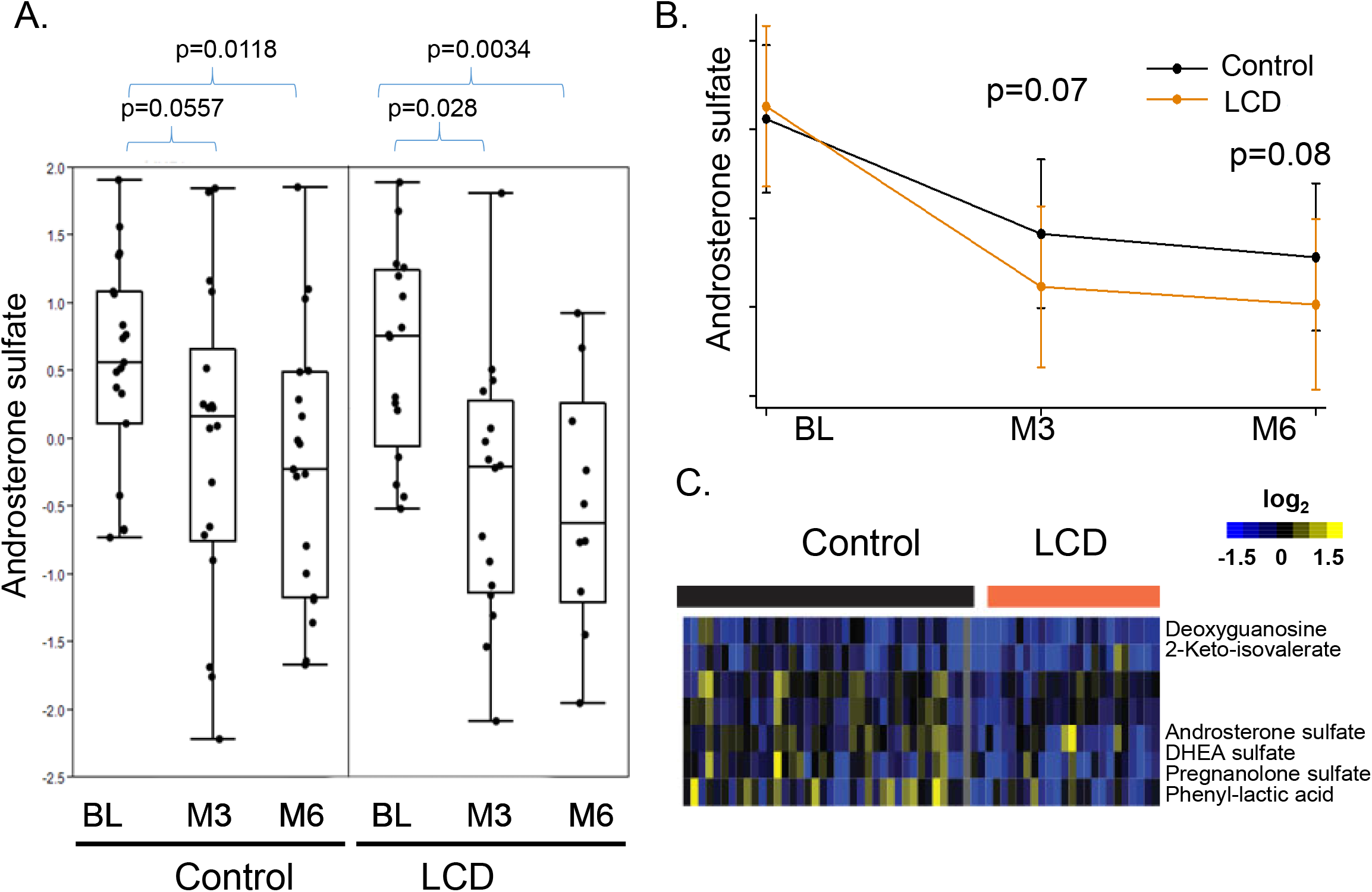
The effects of LCD on the ADT-affected androsterone sulfate and related metabolites. (A) ADT reduced the levels of androsterone sulfate in both control and LCD arms. (B) The mean levels of the androsterone sulfate in the control and LCD arms at the BL, M3 and M6. (C) Heatmap of the clusters of metabolites that were altered in ways similar to androsterone sulfate.

### ADT-induced metabolic changes reversed by LCD

In the control arm, 3-hydroxybutyric acid was reduced at both M3 and M6 (Fig 4A) (21). However, it was significantly increased by ADT at M3 in the LCD arm, and though the increase was somewhat mitigated at M6, it remained higher than BL (Fig 4A). The heatmap also showed that 3-hydroxybutyric acid was reduced in the control arm (blue) and increased in the LCD arm (yellow) (Fig 4B). The opposite ADT-induced changes in control vs. LCD arm of 3-hydroxybutyric acid also co-clustered with similar patterns in 2-hydroxybutyric acid, 3-hydroxy-3-methyl-glutaryl-carnitine and 3-aminobutyrate. Similarly, while many acyl-carnitines were noted to be reduced by ADT in the control arm (21), in LCD arm, ADT paradoxically increased the levels of many acyl-carnitines, including hydroxy-butyryl-carnitine, tiglyl-carnitine, pimelyl-carnitine, 3-hydroxyoctanoyl-carnitine, decanoyl-carnitine, 6-ketodecanoyl-carnitine, dodecanoyl-carnitine, dodecenoyl-carnitine, decanoyl-carnitine, decenoyl-carnitine (Fig 4B). Therefore, the changes of these acyl-carnitines may be tightly associated with the ketone-related metabolites, indicating their potential metabolic connection. Importantly, LCD was able to reverse the reduction of these ketone metabolites and acyl-carnitines as one of the most prominent metabolic effects of ADT.

**Figure 4:**
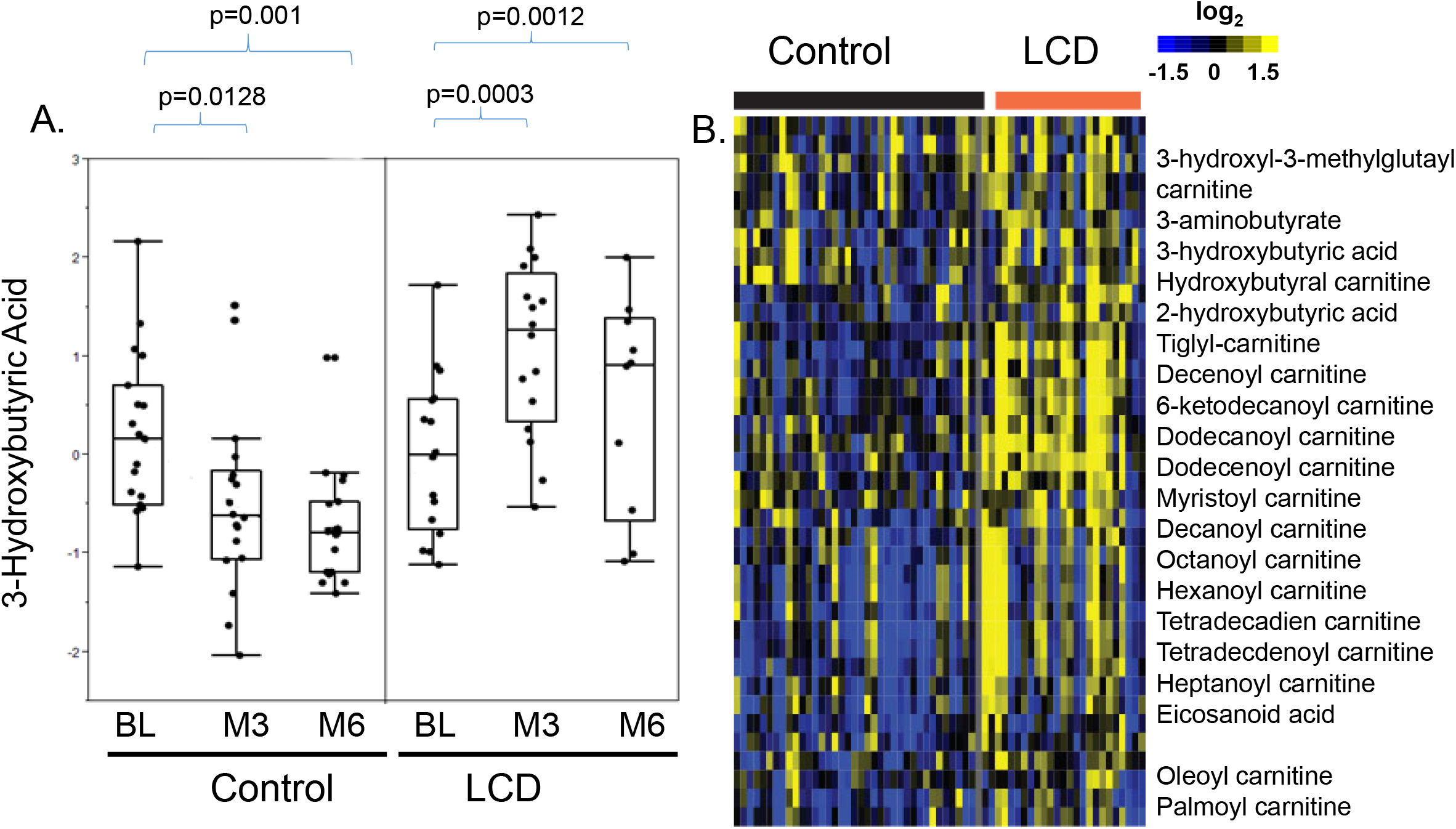
The effects of LCD on the ADT-affected 3-hydroxy-butyric acid. (A) The effects of ADT-induced changes in the 3-hydroxy-butyric acid in the control and LCD arms at M3 and M6. (B) Heatmap of the clusters of metabolites that were altered in ways similar to 3-hydroxy-butyric acid. The statistical significance (p values) of ADT-induced changes of indicated metabolites is indicated.

### Other metabolites whose ADT-induced changes were reversed by LCD

While baseline levels of lactic acid were similar between control and LCD arms, it was slightly increased by ADT in the control arm but reduced in the LCD arm (Fig 5A). This opposite changes lead a significantly higher levels of lactic acids in control than in the LCD arm at both M3 (p<0.0001) and M6 (p=0.0004) (Fig 5A). As shown in the heatmap (Fig 5B), the ADT-induced changes in lactic acid co-clustered with pyruvate, another metabolite in the glycolysis pathway, potentially reflecting glycolysis.

**Figure 5:**
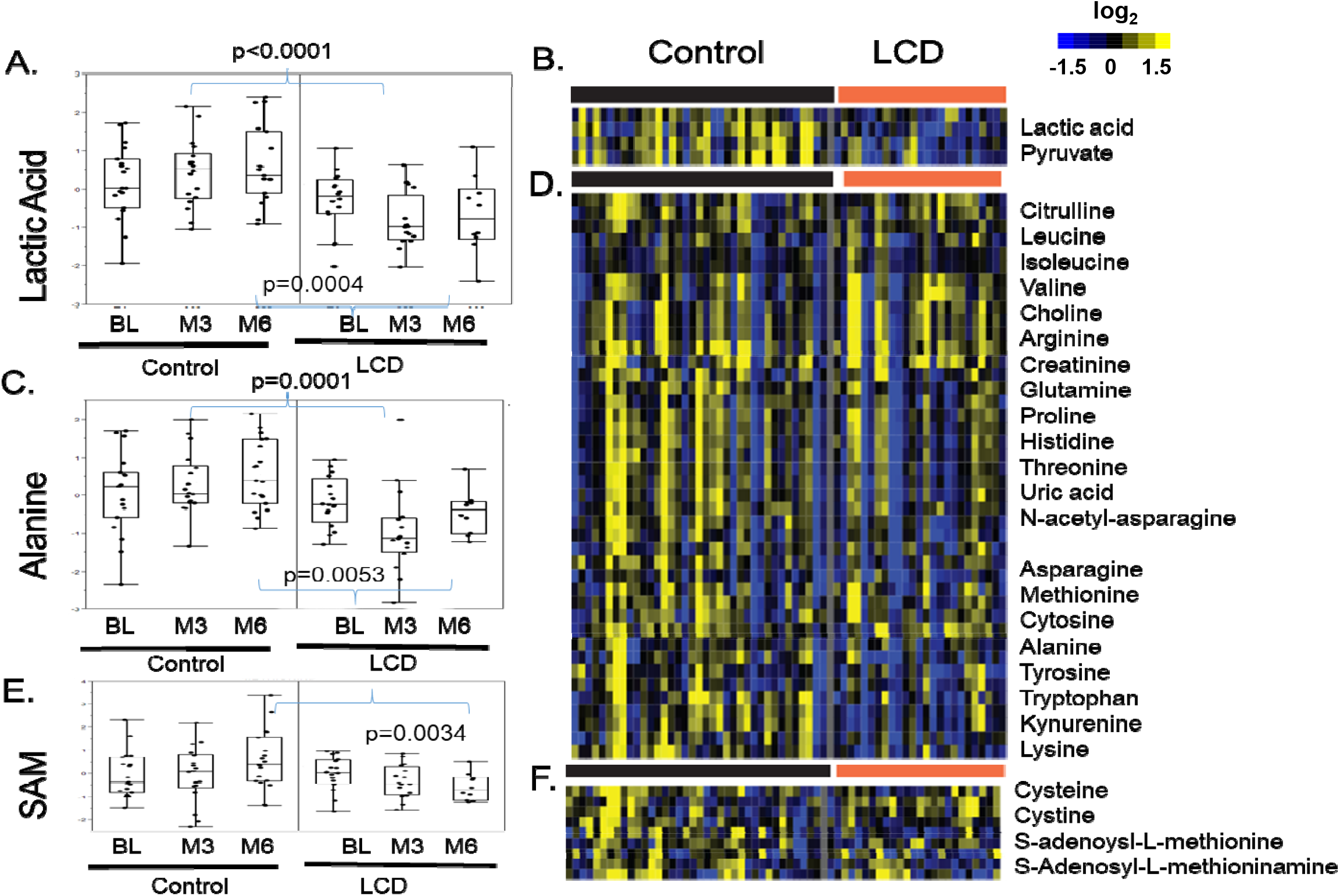
The effects of LCD on the ADT-altered lactic acid, alanine and S-adenosyl-L-methionine (SAM) (A) The effects of ADT-induced changes in the lactic acid in the control and LCD arms at M3 and M6. (B) Heatmap of the clusters of metabolites that were altered in ways similar to lactic acid. (C) The effects of ADT-induced changes in the alanine in the control and LCD arms at M3 and M6. (D) Heatmap of the clusters of metabolites that were altered in ways similar to alanine. (E) The effects of ADT-induced changes in the S-adenosyl-L-methionine (SAM) in the control and LCD arms at M3 and M6. (F) Heatmap of the clusters of metabolites that were altered in ways similar to SAM. The statistical significance (p values) of ADT-induced changes of indicated metabolites is indicated.

Similar trend was also observed for alanine, which remained relatively unchanged by ADT in the control arm, but decreased by ADT in the LCD arm so that the levels were significantly lower in the LCD arm at M3 (p=0.0001) and M6 (p=0.0053) relative to corresponding time points in the control arm (Fig 5C). Such a pattern of ADT-increase that is reversed by LCD was also found for other co-clustered amino acids, including the branched-chain amino acids (valine, isoleucine and leucine) and tyrosine, lysine, leucine, isoleucine, cytosine, methionine, asparagine, histidine, proline, glutamine, and arginine (Fig 5D). There is also a similar reduction in the tryptophan and its metabolite kynurenine in the LCD arm with strong immuno-suppressive function (30).

Similarly, SAM (S-adenosyl-L-methionine) was slightly increased in the control arm and slightly reduced in the LCD arm such that the levels at M6 were significantly different (Fig 5E, p=0.0034). Such a pattern of SAM was found to be co-clustered with several other sulfur-containing amino acids, including S-adenosyl-L-methioninamine, thymine, cysteine and cystine (Fig 5F). The pattern of changes in control and LCD arm was also similar to that of other amino acids (Fig 5D). Finally, we also noted that ADT increased the levels of succinyl-adenosine at M6 in the control arm (Supplemental figure 1), which was abolished by LCD.

### Correlation of blood chemistry with the ADT-induced changes in androsterone sulfate

Although all androgens were expected to be reduced by ADT in both arms, androsterone sulfate was the most consistently reduced metabolite at both time points in both arms (Fig 1A, C). However, there were still individual heterogeneity in the ADT response. Therefore, we reasoned that ADT-reduced androsterone sulfate may provide a quantitative assessment ADT response and quantify the correlation with different blood chemistry and serum metabolomic. First, we calculated various blood chemistry with androsterone sulfate. In the control arm, the ADT-induced androsterone sulfate was negatively correlated with glucose levels at both M3 (p<0.001, r=-0.811) and M6 (p=0.0021, r=-0.660) (Fig 6A, B). This indicates that a stronger reduction of androsterone sulfate is associated with increased glucose levels, supporting the diabetogenic tendency of the ADT. In contrast, the changes in androsterone sulfate were not significantly correlated with total cholesterol, LDL or insulin in either control or LCD arm (Supplemental figure 2A-C). Importantly, these correlations between glucose and androsterone sulfate disappeared in the LCD arm, suggesting that LCD disrupted the tight correlation between ADT response and increased serum glucose (Fig 6A, B).

**Figure 6:**
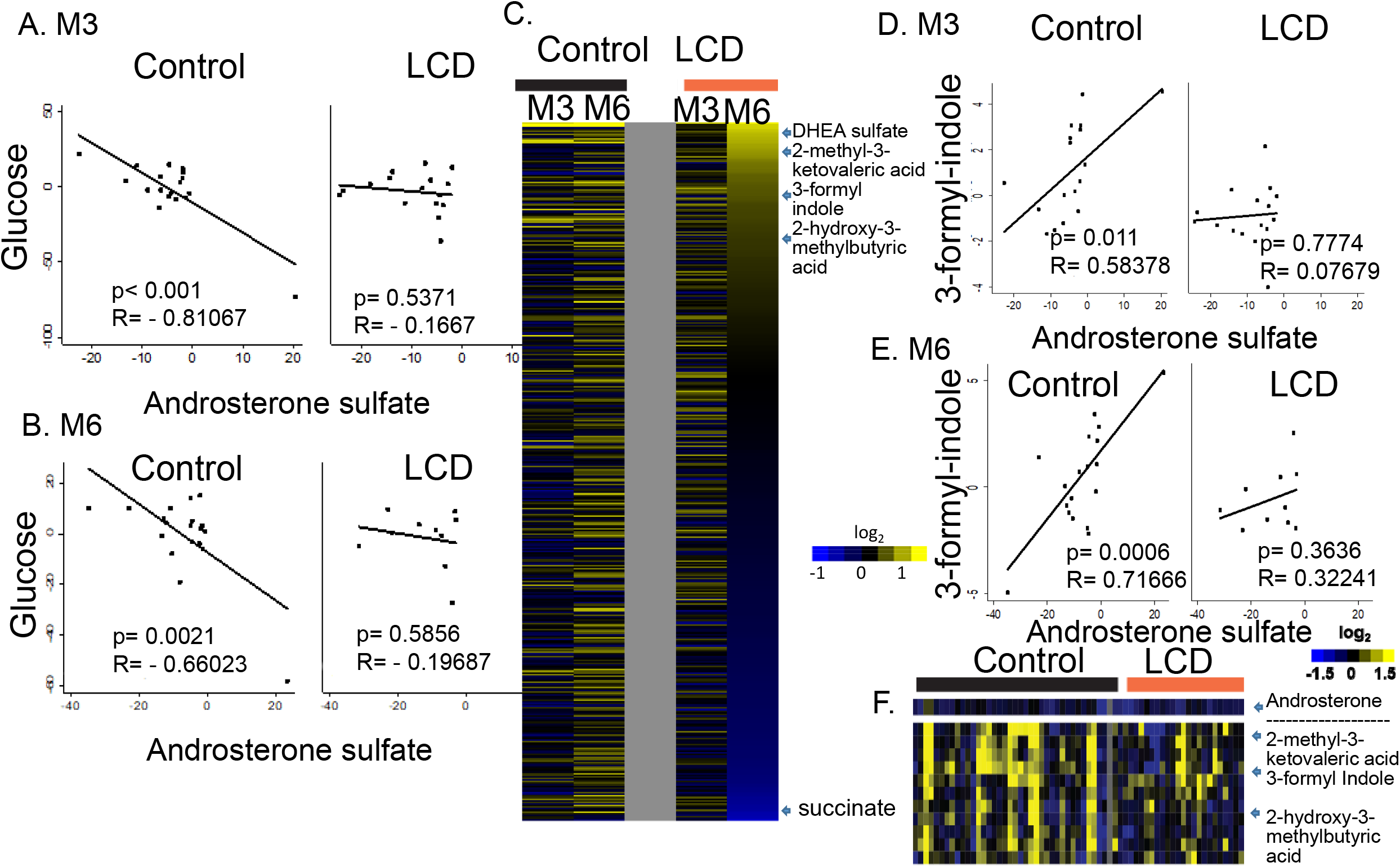
Serum chemistry and metabolites correlated with ADT-reduced androsterone sulfate. (A-B) The correlation between the ADT-induced changes in the androsterone sulfate and serum glucose in the control and LCD arm after 3 and 6 months (B) of ADT. (C) Heatmap of the correlation between the ADT-induced changes in metabolites with androsterone sulfate. Yellow: Positive correlation and Blue: Negative correlation. The metabolites showing the most positive and negative with the selected metabolites indicated. (D, E) The correlation plots between the ADT-induced changes in the 3-formyl-indole with androsterone sulfate at M3 (D) and M6 (E) in the control and LCD arms. (F) The effects of ADT on the 3-formyl indole and other indicated co-clustered metabolites.

### Identify serum metabolites whose changes correlate with androsterone sulfate

Next, we calculated the ADT-induced changes in serum metabolome whose correlated with the ADT-affected androsterone sulfate in all groups and ranked M6-LCD metabolites from the highest (close to +1) to the lowest correlation (close to −1) in a heatmap (Fig 6C, complete list in the Supplemental Table 3). Among the metabolites positively correlated metabolites in LCD at M6, many metabolites were also positively correlated (yellow in the heatmap) in the control arm at both M3 or M6, but not in the LCD at M3 (black/blue in the heatmap) (Fig 6C). Therefore, the LCD altered many observed correlations in the control arm at M3-LCD, but this was partially restored at M6-LCD. For example, there is a strong correlation between androsterone sulfate and other androgens, including pregnenolone sulfate and DHEA-androsterone in the control arm at both M3 and M6 (Fig 6C) (21). While such correlation still existed between pregnenolone and androsterone (Supplemental figure 3A), LCD disrupted the DHEA-androsterone correlation at M3 (Supplemental figure 2B), which was partially restored at M6 (Supplemental figure 3B). In addition, there was a positive correlation between androsterone sulfate and 2-methyl-3-ketovaleric acid and 2-hydroxy-3-methylbutyric acid as well as 3-formyl-indole. 3-formyl-indole is known as Indole-3-carboxaldehyde (I3A) or indole-3-aldehyde as a metabolite of dietary L-tryptophan that is synthesized by the gut microbiome. There was a strong correlation with 3-formyl-indole at both M3 (p=0.011) and M6 (p<0.001) in the control arm (Fig 6D). This relationship disappeared in the LCD arm at M3 and resumed slightly at M6 (Fig 6E). Therefore, LCD disrupted the correlative relationship between androsterone sulfate with many metabolites at M3, which were partially restored at M6. In contrast, the metabolites which were negatively correlated with the androsterone sulfate in LCD at M6 were only seen in the LCD arm, not in the control arm. In fact, the correlation was the strongest in the LCD at M6 (Fig 6C). For example, succinate was negatively correlated with androsterone sulfate, indicating that the degree of its increase was associated with the degree of ADT-reduced androsterone sulfate. Next, we examined how ADT affected the 3-formyl-indole in the control and LCD arms in the heatmap. We found that 3-formyl-indole was co-clustered with 2-methyl-3-ketovaleric acid and 2-hydroxy-3-methylbutyric acid which also showed strong positive correlation with androsterone (Fig 6F). Interestingly, we noted that ADT mostly increased 3-formyl-indole as well as 2-methyl-3-ketovaleric acid and 2-hydroxy-3-methylbutyric acid in the control arm (Fig 6F). In the control arm, 3-formyl-indole in was increased in ∼60 % participants and reduced in ∼40% participants. Interestingly, the participants with an increase in the 3-formyl-indole had less androsterone sulfate reduction than the participants with reduced 3-formyl-indole (Fig 6D, E, F), thus accounting for the positive correlation between these two metabolites. In the LCD arms, the changes of 3-formyl-indole in response to ADT were dampened and the correlations between 3-formyl-indole and androsterone vanished.

Together, these data indicate that LCD at M3 disrupted many correlations between ADT-affected androsterone with glucose, DHEA, 3-formyl-indole seen at both M3 and M6 in the control arms. However, these correlations were partially restored at M6 in LCD arms, suggesting metabolic adaptations.

## Discussion

In this study, we applied MS-based metabolomic profiling to determine the effects of LCD after 3 and 6 months of ADT on the serum metabolome of PC participants. The key metabolic changes observed as a result of ADT can be summarized into five areas: 1) steroid metabolism, 2) ketogenesis, 3) acyl-carnitines metabolism, 4) glucose and glycolysis, and 5) microbiome-associated metabolites. These analyses showed that LCD not only did not negate the effect of ADT to reduce androgens but may slightly enhance this effect although these results did not reach statistical significance. LCD also reversed ADT-reduced ketogenesis and acyl-carnitines. LCD also disrupted the correlation between high serum glucose and lower androsterone sulfate and reversed the ADT-induced changes in glycolysis and amino acid metabolisms. Together, LCD significantly reversed many of the ADT-impacted metabolic changes in serum and therefore may mitigate the metabolic side effects of ADT, such as reduced weight, improved insulin resistance, hemoglobin A1c, HDL, and triglyceride as previously reported in our CAPS1 paper(10). Furthermore, some of these LCD-induced metabolomic changes may suggest that LCD can be used as adjuvant therapy that may contribute to tumor-suppressing effects via different mechanisms, though this would require formal testing in a large and longer study. These potential tumor-suppressing metabolic changes of LCD include the possible enhancement of androgen suppression, reduced tumor glycolysis and lower immune-suppressing metabolites (lactate and kynurenine). Although the current study is under-powered to establish these benefits, our results may support future large clinical trials of LCD with ADT since LCD has been shown to be safe and well-tolerated. Consistently, a follow-up CAPS2 study has shown LCD is safe, does not negatively slow tumor growth and may suggestively delay tumor growth as measured by slower PSA doubling times among men with recurrent PC (31). Furthermore, ketogenic diets also have significant anti-tumor effects the murine models (32). Whether an LCD can slow tumor growth or not, our results support the potential for LCD to prevent ADT-induced metabolic effects. However, some metabolic changes of LCD were transient and seen only at M3, suggesting potential metabolic adaptation that may have reduced the impact of LCD at M6.

To the best of our knowledge, this the first study to employ serum metabolomics to investigate the effects of LCD in the context of ADT. Several previous studies have investigated the metabolic response to LCD in the contexts of other cancers. For example, LCD in people with pancreatic cancer increased serum β-hydroxybutyrate and total ketone levels as well as the glycerophospholipid and sphingolipid metabolisms (33). In another study of mouse xenografts treated with LCD, breast cancer leads to serum metabolic changes, which were reversed with LCD. LCD also affected the amino acid metabolism and fatty acid transport, suggesting a tumor-suppressing mechanism of LCD when combined with tumor therapeutics (34). Our data add to the expanding literature on the metabolomic effects of LCD in the contexts of ADT treatments for PC.

In the primary paper describing the clinical outcomes of CAPS1 trial, LCD did not significantly affect PSA compared with the control arm (10). However, as mentioned earlier, a follow-up randomized CAPS2 trial, LCD, has been show to delay PSA doubling times (31). Here, we found that androsterone sulfate was the most consistently ADT-reduced androgen in all participants, supporting it as the best biomarker in this study to quantify the efficacy of ADT. In contrast, other androgens may be affected by LCD and not as reliable. Interestingly, while ADT reduced androsterone sulfate in both arms, the reduction showed a tendency to be even more pronounced in the LCD arm, consistent with the longer PSA doubling times in CAPS2 (31). Future research with more participants is warranted to determine whether LCD may enhance the ADT-reduced androsterone sulfate or not. Moreover, given that lower nadir androgen values on ADT are associated with reduced risk of progression (35), this raises the possibility than a LCD may enhance the anti-PC effects of ADT, though this requires formal testing in future studies.

In CAPS1, ketone bodies were shown to be decreased by ADT in the control arm (21). Ketone bodies are also postulated to affect insulin resistance and increased ketone bodies during ketogenic diets may account for the reduced insulin resistance (36). Therefore, ADT-associated reduced ketone bodies may contribute to ADT-induced insulin resistance and justify the potential of ketogenic LCD to reverse these effects. As expected, we noted that LCD increased 3-hydroxybutyric acid at both time points compared to baseline (Fig 2, 3). These results suggest that LCD intervention was effective in overcoming ADT’s impact in reducing this metabolite, maybe through the activation of fatty acid β-oxidation and ketogenesis. The increase of 3-hydroxybutyric acid was less dramatic at M6 compared to M3, which may be due to a possible metabolic adaptation or lower adherence to the LCD. However, since the carbohydrate intake and adherence to the LCD were similar at M3 (carb intake: 74.4 gm) and M6 (78.9 gm) vs. BL (227.2 gm) (10), we speculate results are more likely due to adaptation. However, these results could also mean an increased conversion of 3-hydroxybutyric acid to acetyl-CoA and consumption as an alternative energy source. Elevated levels of this ketone body may be associated with low blood glucose in response to the LCD and disruption between the androsterone sulfate and glucose increase (Fig 6A, B). Future research is warranted to clarify the potential of LCD in reducing ADT-induced insulin resistance and to examine how to sustain this potential long-term.

Acyl-carnitines are generated by carnitine acyltransferases from combining carnitine and acyl-CoA as metabolic intermediates of fatty acid metabolism critical for the prostate cancers(37,38). Recent findings have suggested that the carnitine system could serve as a reservoir to finely trigger the metabolic flexibility of cancer cells (39) involved in the bi-directional transport of acyl moieties between cytosol to mitochondria in tuning the switch between glucose and fatty acid metabolism. High levels of acyl-carnitines are found in diet-induced obesity and insulin resistance in human and mouse models(40,41). Therefore, elevated acylcarnitine has been postulated to contribute to insulin resistance phenotypes (42) of ADT. We found that LCD reversed the ADT-reduced acyl-carnitines and the increased acyl-carnitines were strongly clustered with increased ketone-related metabolites. Together, these findings suggest a metabolic connection between ketogenesis and changes of acyl-carnitines, indicating the ability of LCD to reverse these metabolic changes in response to ADT.

Our results also show that while ADT slightly increased lactic acidic and pyruvate, LCD significantly lowered these glycolysis-related metabolites. Warburg effects and increased glycolysis are one of the major hallmarks of cancer. Therefore, the slight increase in the lactic acidosis by ADT suggests an increase in glycolysis during ADT. However, the restriction of glucose and other changes in the LCD arm may contribute to the reduced glycolysis and lactic acid production, which may contribute to the anti-tumor efficacy. Recently, lactic acidosis has been shown to affect tumor gene expression, select tumor mutations (43-45) and suppress tumor immune microenvironments (46,47). Therefore, the reduction of lactic acid by LCD may also mitigate the immune suppression environments that can promote tumor growth. Similarly, LCD also reduced the levels of kynurenine (Fig 5D), another immune-suppressive metabolite (48) associated with tumor progression. Therefore, the ability of LCD to reduce both lactate and kynurenine may suggest the potential of LCD to overcome immune suppression. A recent study showed that low protein diets, but not LCD, enhance immune surveillance and reduce tumor growth (49). Our findings may suggest that LCD may have a better immuno-modulatory function in the contexts of ADT for prostate cancer.

Previously, we found that the degree of ADT-induced androsterone sulfate was strongly associated with the changes in indole-3-carboxaldehyde (ICA), a microbiota-derived metabolite that stimulates aryl hydrocarbon receptors in intestinal immune cells that produce IL-22 during mucosal reactivity and inflammation in mice (22). ICA treatments reduced gut inflammation, limited epithelial damage and reduced transepithelial bacterial translocation, and decreased inflammatory cytokines (50). Consistently, ADT increased the levels of ICA (Fig 6F), which may contribute to the reduced inflammatory bowel diseases (50). Since LCD mitigates the ADT-induced changes of ICA, we expect that LCD may also reduce the IBD protection effects of ADT. Recent studies showed an intense chemical exchange between host and microbiome during ketogenic diets (51). Ketogenic diet-induced increases in ketone bodies reproducibly inhibited bifidobacterial growth and reduced intestinal pro-inflammatory Th17 cells (51). These temporary effects indicate that many of the diet-induced changes in the microbiomes may be compensated by the hosts’ adaptive response. Therefore, it will be important to identify the nature and mechanisms of these metabolic adaptation to sustain the metabolic effects and therapeutic benefits of LCD.

However, it is important to point out that our results should be interpreted within the limitations of our study. A small sample size of the cohort may limit our power to detect subtle differences that may be of importance. For example, the potential of LCD to enhance ADT-reduced androsterone sulfate should be tested in a larger cohort to define whether this is a reproducible finding. Future studies are needed to test the potential of combining adjuvant LCD to enhance the efficacy of ADT. In addition, future studies are needed to correlate these metabolomic changes with the long-term outcomes including tumor control and metabolic side effects of ADT such as diabetes risk.

## Conclusion

Our findings indicate the metabolic basis for the potential of LCDs in reversing many of the side effects of ADT. However, some of the reversed effects appeared temporary as they were less pronounced at 6-month than at 3-month. Future studies are needed to understand how the effects can be sustained. Further, our study suggests that LCDs may enhance the effect of ADT by reducing androsterone sulfate and other metabolic mechanisms, though future research is needed to confirm this finding and understand the underlying mechanisms.

## Data Availability Statement

The metabolomic data will be made available to the academic community upon publication of the manuscript.

## Funding and Acknowledgement

American Urological Association Foundation, BERG, NIH, and Robert C. and Veronica Atkins Foundation

## Conflict of interest statement

No conflict of Interest.

## Figure legends

**Supplemental Figure 1: The effects of LCD on the ADT-affected succinyl-adenosine** The effects of ADT-induced changes in the succinyl-adenosine in the control and LCD arms at M3 and M6.

**Supplemental Figure 2** (A-C) The correlation between indicated ADT-induced androsterone sulfate with the cholesterol (A), LDL (B) and insulin (C) in the control and LCD arm at M3 and M6.

**Supplemental Figure 3** (A-B) The correlation between indicated male hormones with the ADT-induced changes in the androsterone sulfate in the control and LCD arm at M3 and M6.

**Supplemental Table 1**: Top metabolites affected by the ADT in the LCD arm at M3

**Supplemental Table 2**: Top metabolites affected by the ADT in the LCD arm at M6

**Supplemental Table 3**: The serum metabolites ranked by the correlation with the ADT-induced changes in androsterone sulfate.

Supplemental file: Python script to derive the changes of metabolites in response to ADT.

## Notes

### Competing Interest Statement

The authors have declared no competing interest.

