## Supplementary_Figures for "The effects of low-carbohydrate diets on the metabolic response to androgen-deprivation therapy in prostate cancer"

Supplemental Figure 1

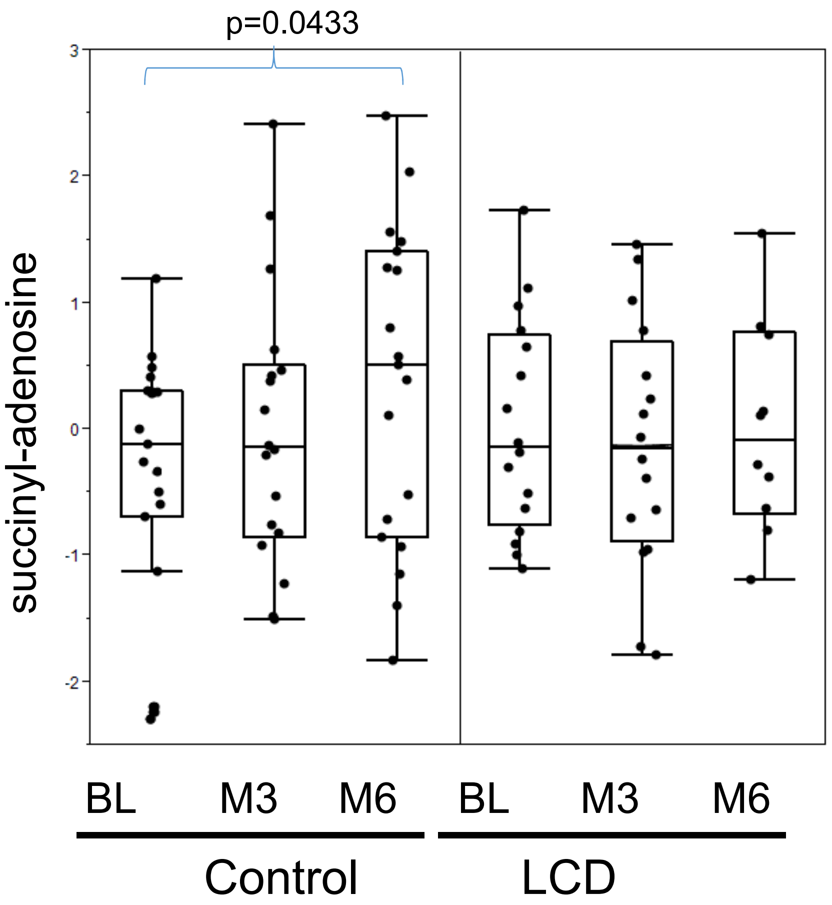


**Supplemental Figure 1: The effects of LCD on the ADT-affected succinyl-adenosine**

(A) The effects of ADT-induced changes in the succinyl-adenosine in the control and LCD arms at M3 and M6.

Supplemental Figure 2A

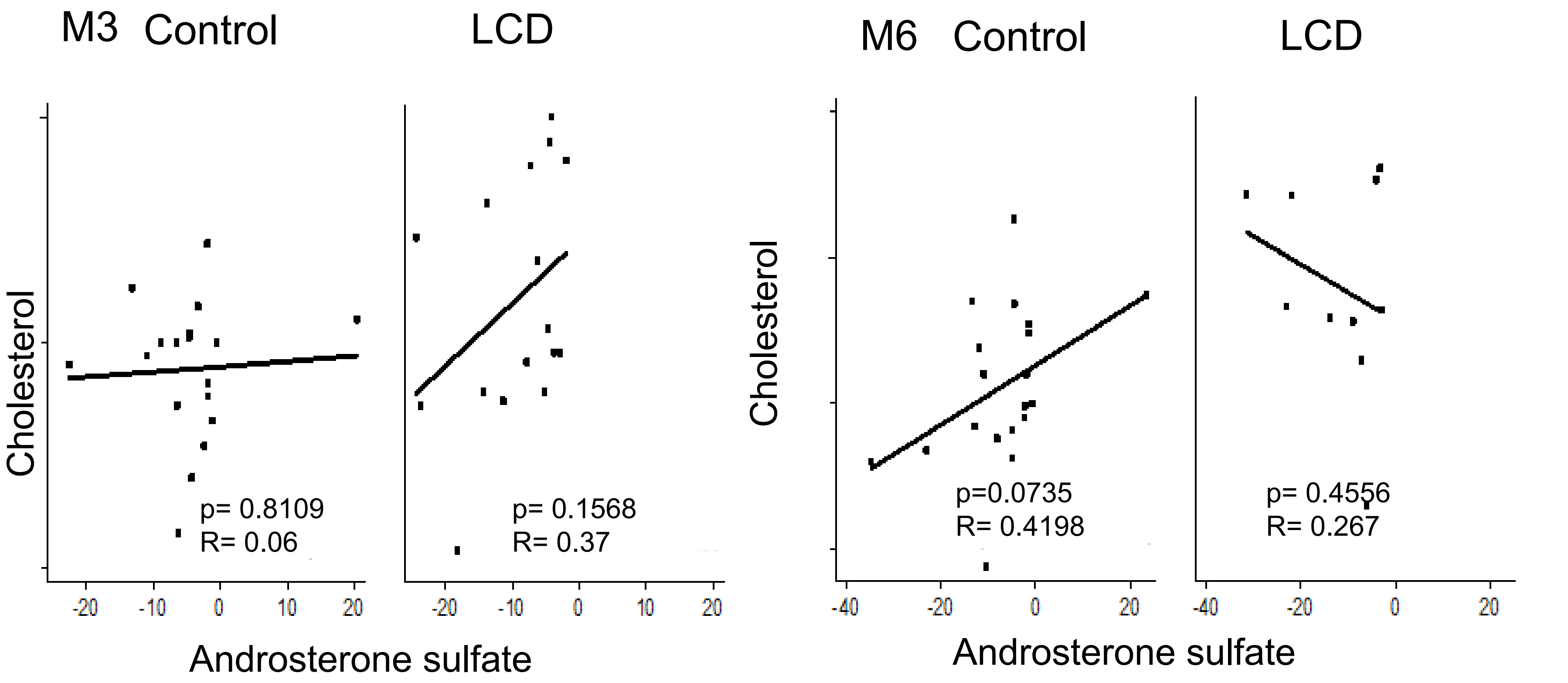


**Supplemental Figure 2** (A-C) The correlation between indicated ADT-induced androsterone sulfate with the cholesterol (A), LDL (B) and insulin (C) in the control and LCD arm at M3 and M6.

Supplemental Figure 2B

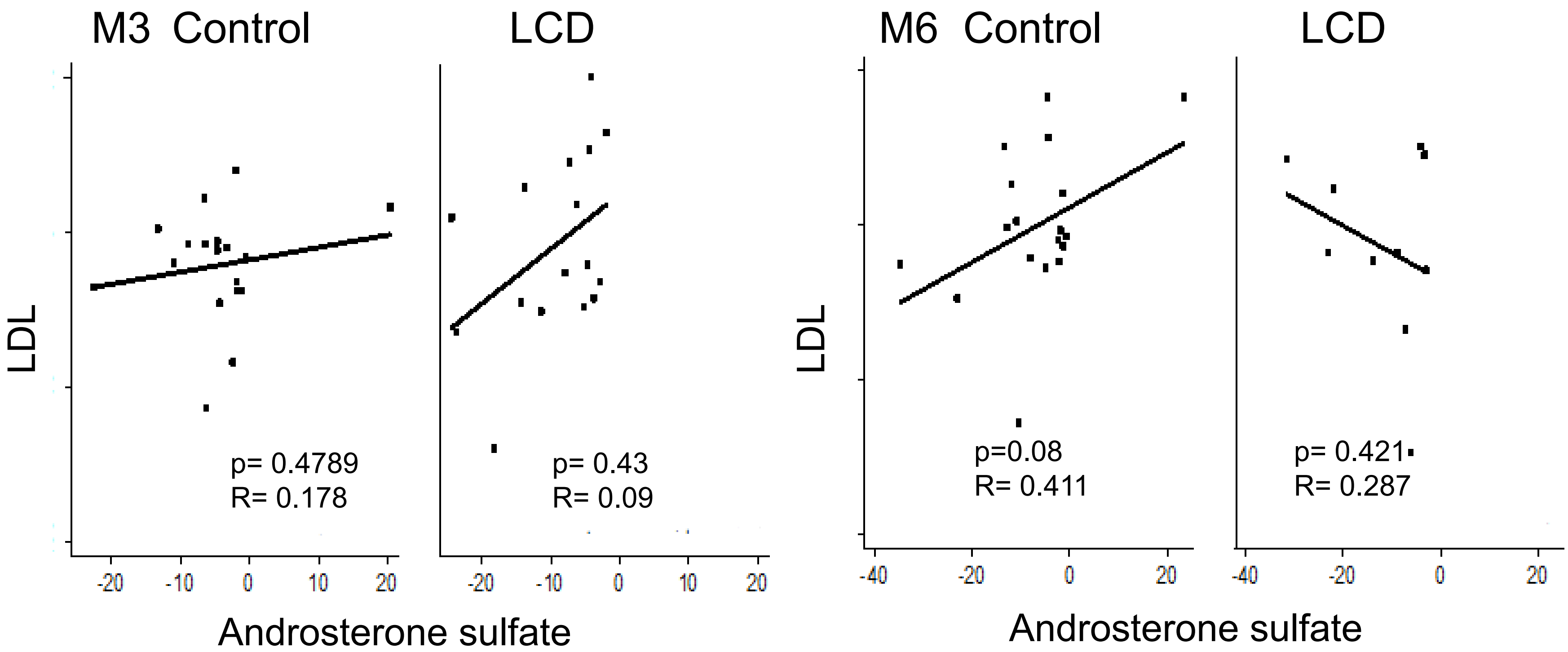


**Supplemental Figure 2** (A-C) The correlation between indicated ADT-induced androsterone sulfate with the cholesterol (A), LDL (B) and insulin (C) in the control and LCD arm at M3 and M6.

Supplemental Figure 2C

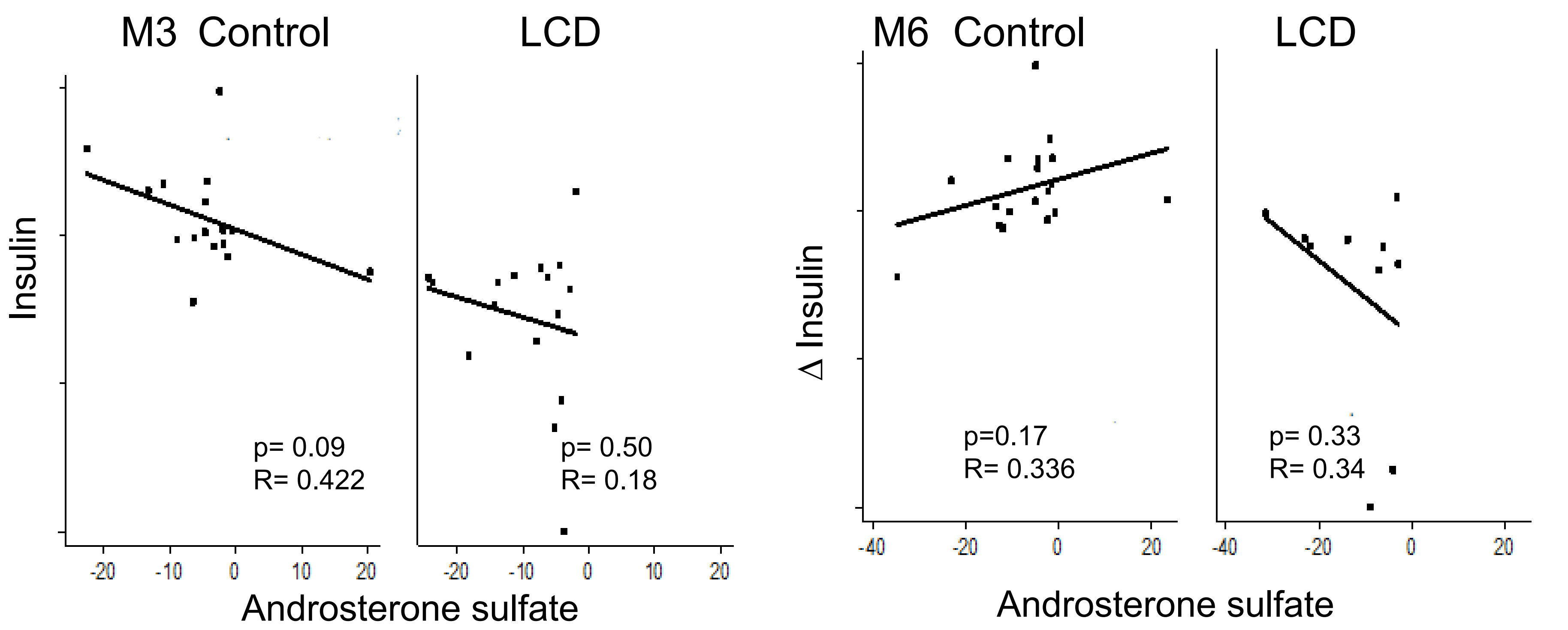


**Supplemental Figure 2** (A-C) The correlation between indicated ADT-induced androsterone sulfate with the cholesterol (A), LDL (B) and insulin (C) in the control and LCD arm at M3 and M6.

Supplemental Figure 3A

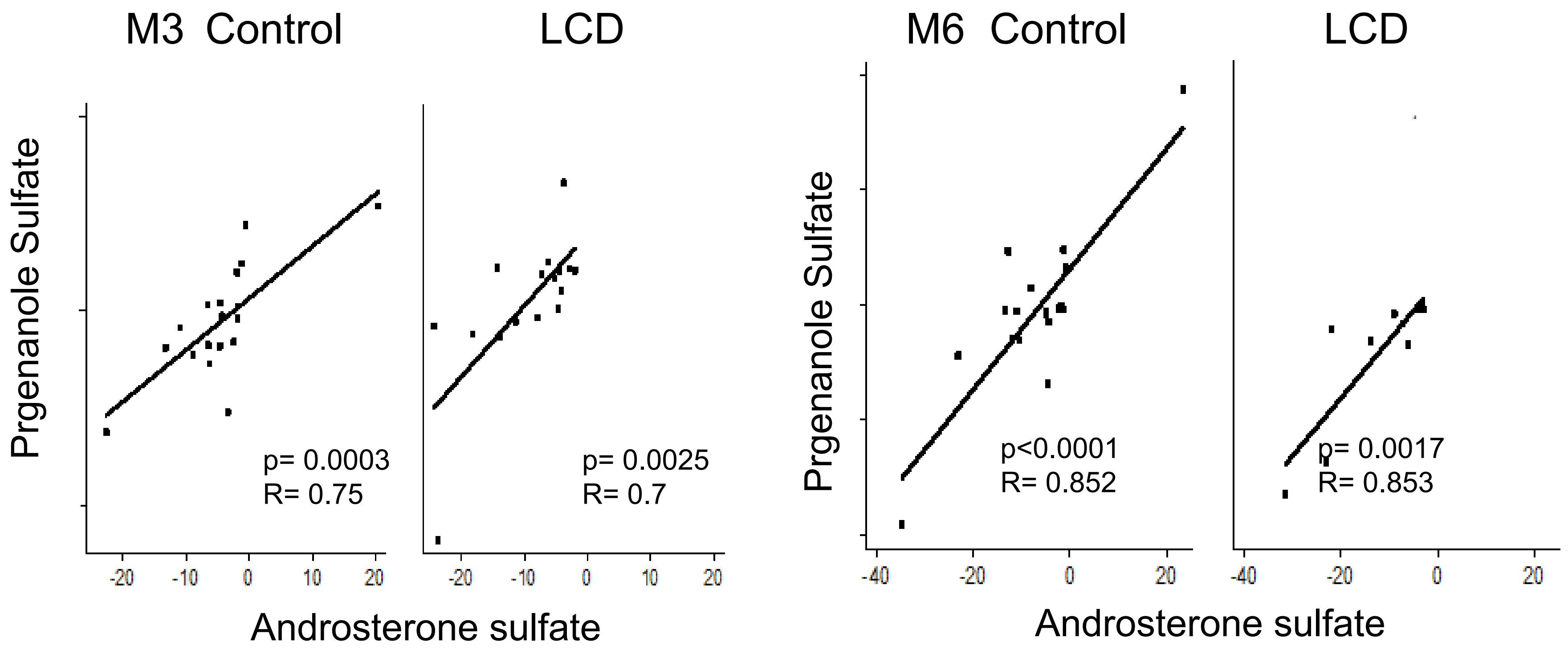


**Supplemental Figure 3** (A-B) The correlation between indicated male hormones with the ADT-induced changes in the androsterone sulfate in the control and LCD arm at M3 and M6.

Supplemental Figure 3B

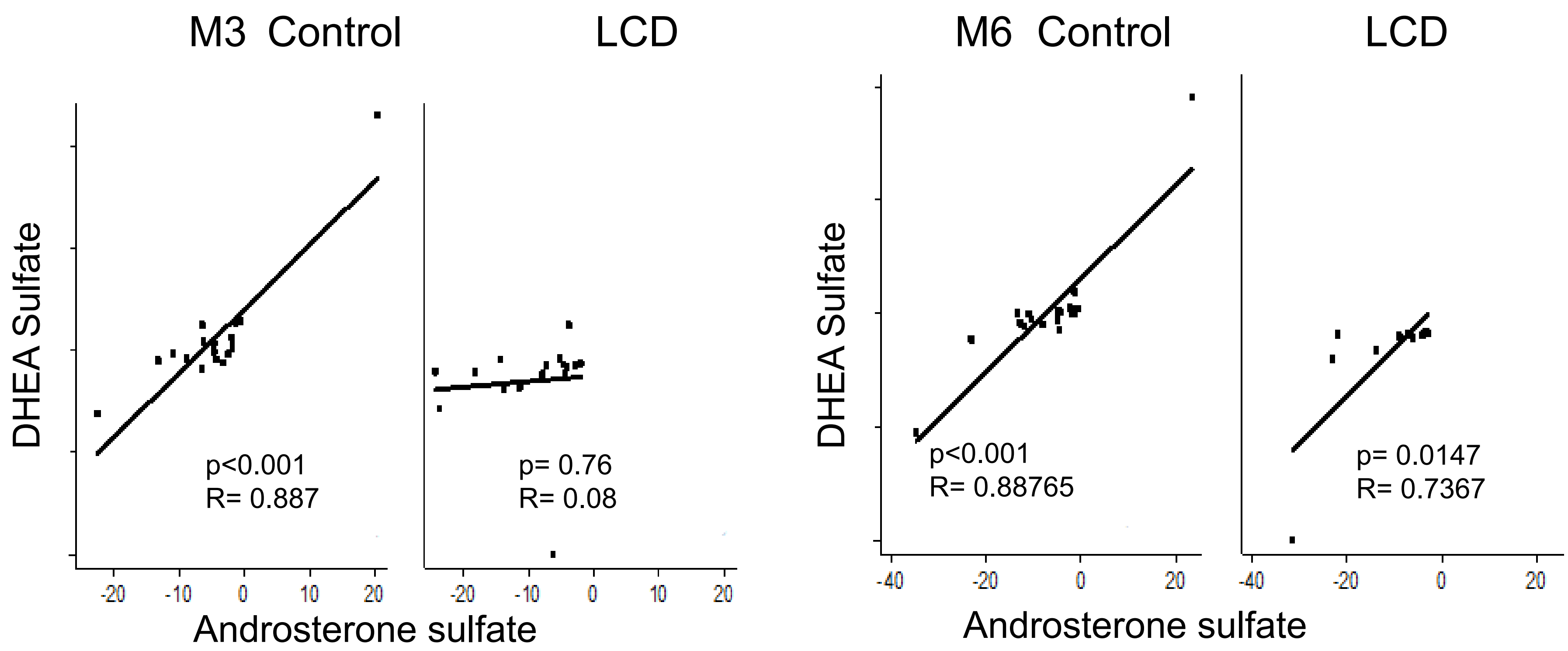


**Supplemental Figure 3** (A-B) The correlation between indicated male hormones with the ADT-induced changes in the androsterone sulfate in the control and LCD arm at M3 and M6.
